# Using Unstructured Crowd-sourced Data to Evaluate Urban Tolerance of Terrestrial Native Species within a California Mega-City

**DOI:** 10.1101/2023.12.05.570260

**Authors:** Joseph N. Curti, Michelle Barton, Rhay G. Flores, Maren Lechner, Alison Lipman, Graham A. Montgomery, Albert Y. Park, Kirstin Rochel, Morgan W. Tingley

## Abstract

In response to biodiversity loss and biotic community homogenization in urbanized landscapes, City managers around the world are increasingly working to conserve and increase urban biodiversity. Accordingly, around the world, previously extirpated species are (re)colonizing and otherwise infiltrating urban landscapes, while once abundant species are in various states of decline. Tracking the occurrence of traditionally urban intolerant species and loss of traditionally urban tolerant species should be a management goal of urban areas, but we generally lack tools to study this phenomenon. To address this, we first used species’ occurrences from iNaturalist, a large collaborative dataset of species observations, to measure an urban association index (UAI) for 967 native animal species that occur in the city of Los Angeles. On average, the occurrence of native species was negatively associated with our composite measure of urban intensity, with the exception of snails and slugs, which instead occur more frequently in areas of increased urban intensity. Next, we assessed 8,348 0.25 x 0.25 mile grids across the City of Los Angeles to determine the average grid-level UAI scores (i.e., a summary of the UAIs present in a grid cell, which we term Community Urban Tolerance Index or CUTI). We found that areas of higher urban intensity host more urban tolerant species, but also that taxonomic groups differ in their aggregate tolerance of urban areas, and that spatial patterns of tolerance vary between groups (e.g., mammals are not the same as birds). The framework established here was designed to be iteratively reevaluated by city managers of Los Angeles in order to track the progress of initiatives to preserve and encourage urban biodiversity, but can be rescaled to sample different regions within the city or different cities altogether to provide a valuable tool for city managers globally.

## INTRODUCTION

The Earth is experiencing an extinction crisis, with modern species extinction rates, based on vertebrate taxa, estimated to exceed background rates of extinction by at least an order of magnitude (Ceballos et al. 2015; McCallum 2015). In this contemporary era of species loss, there are a multitude of factors driving global declines including habitat loss, invasive species, disease, direct exploitation, pollution, and human-caused climate change (Parmesan and Yohe 2003; Sodhi et al. 2008; Szabo et al. 2012; IPBES 2019; Munstermann et al. 2022). Many of the effects of these extinction drivers are increased due to synergistic interactions (Brook et al. 2008); particularly, urbanization is well known to compound all of these drivers of extinction (Brook et al. 2008; Fenoglio et al. 2021; Ruas et al. 2022). Globally, urban cover is predicted to increase by 2.5% between 2000 and 2030, such that urbanization will continue to increase as a driver of biodiversity loss (Seto et al. 2012). Increases in urban cover are predicted to grow especially fast within global biodiversity hotspots, potentially by 200% between 2000 and 2030 (Seto et al. 2012), which could further exacerbate rates of species decline. With this predicted increase in urban cover, city managers and conservation biologists can work collaboratively to make cities more hospitable to native biodiversity in order to help avert increasing levels of extinctions.

The correlation between urbanization and biodiversity loss is well documented, especially in birds (Batáry et al. 2018), arthropods (Faeth et al. 2011; Fenoglio et al. 2020), and plants (Rega-Brodsky et al. 2022). Research in urban biodiversity has traditionally focused on quantifying changes in species richness along the urban-rural gradient (e.g., McDonnell and Picket 1990), especially on native bird populations. Studies demonstrate that native woodland bird species tend to be replaced by urban-adapted species in more urbanized habitats nearer to the city center (Blair 2004; Blair and Johnson 2008). This pattern appears to be stronger for some groups of birds. For example, migratory species have been found to be more sensitive to urbanization than resident species (Husté and Boulinier et al. 2007). Patterns of bird diversity are also tied to factors such as vegetation community structure. For example, canopy cover and native plant species diversity correlate with increases in native bird species richness in urban woodlands (Fontana et al. 2011; de Toledo et al. 2012; Wood and Esaian 2020); whereas, increases in lawn cover are related to increases in non-native and synanthropic species richness (Paker et al. 2014). These studies demonstrate that the mechanisms underlying urban biodiversity are complex, and the narrow geographic and taxonomic breadths of current studies make it difficult to generalize patterns in ways that can help cities improve their biodiversity (Faeth et al. 2011; Magle et al. 2012; Beninde et al. 2015; Rega-Brodsky et al. 2022).

Considering the projected increase in urban land cover within the next decade, the future of urban biodiversity will ultimately rely on the ability of global cities to attract and maintain populations of species that are largely considered urban intolerant. Well planned cities can preserve and restore the habitat requirements of native species including heterogeneous landscapes, migratory stopover sites, and increased gene flow in some species (Spotswood et al. 2021). Likely as a result of these benefits, some species’ ranges have increased in urban areas over the past century (Wehtje 2003; Veech et al. 2011; Ancilloto et al. 2016; Urošević et al. 2016). Some studies have examined what factors increase and maintain urban biodiversity, for example, by evaluating the minimum number of native trees in urban residential yards needed to maintain diverse bird communities (Lerman and Warren 2011) or quantifying native species gained in planted rooftop gardens (Wooster et al. 2022). While these projects can help inform policy geared towards supporting and enhancing urban biodiversity, city managers still lack a comprehensive tool that can track spatio-temporal changes in urban biodiversity at the community level. As more native species are threatened with extirpation, and some expand their ranges into urban environments, creating a tool that can track changes in urban diversity and community composition is more important than ever before.

In order to monitor diversity patterns and quantify the effects of varying levels of urbanization on different groups of native species, large amounts of data are needed across multiple taxonomic groups and across broad areas of the urban environment. To correct for researcher biases that lead to datasets with limited geographic scope and taxonomic coverage, many studies have turned to large crowd-sourced datasets (Yang 2020; Uchida et al. 2021; Rega-Brodsky et al. 2022). Such datasets are often referred to as ‘unstructured’ in that there is no required protocol for data collection, resulting in data that vary widely in their quality, organization, and information content (Gandomi and Haider 2015). One such platform, iNaturalist, has over 74 million observations for over 342,000 different species globally, 58% of which come from developed (i.e. urbanized) areas (Di Cecco et al. 2021). The size of this dataset gives it great potential for tracking and managing urban biodiversity. For example, Callaghan et al. (2020) used community science data from metropolitan regions across Boston to quantify biodiversity responses at the species and community levels to gradients of different urbanization variables across the landscape. These data were at a small-enough spatial unit to influence local policymaking. Large-scale public participatory datasets make urban biodiversity assessments at large spatial scales possible, even in cities, which tend to contain private lands that are largely excluded from structured biodiversity surveys (Li et al. 2019).

Despite the abundance of data points from programs like iNaturalist, there are challenges associated with using these unstructured datasets to measure and manage urban biodiversity. For example, opportunistic sampling may lead to biases in data, as sampling effort is not equal across space, time, and taxonomic groups (Kamp et al. 2016; Rapacciuolo et al. 2021), potentially causing differences in user methodology to be misinterpreted as temporal or spatial changes in populations (Bayraktarov et al. 2019). However, there has been much work to account for and help mitigate these inherent biases in unstructured data, including using both models and data processing to better account for unequal observations across space and time (Van Strien et al. 2013; Isaac et al. 2014; Steen et al. 2019). Higher order taxa may also be used as indicators instead of single species to reduce taxonomic bias and differences in detectability for any given species (Rapacciuolo et al. 2021).

Here, we describe and implement an approach to spatially and temporally characterize urban tolerance of native species within the city of Los Angeles, California, USA using unstructured species occurrence data from iNaturalist. This approach was initially conceived to support the LA Biodiversity Index Baseline Report published by the Los Angeles Department of Sanitation and Environment (LASAN 2022a) through the creation of an evaluative metric (Metric 1.2b; LASAN 2018, 2020) that represents and monitors “Native Species Presence in Urban Areas.” We refer to this index as a “Community Urban Tolerance Index” (CUTI), as it broadly aims to track how well native species that are often urban intolerant occur within Los Angeles by rating spatial units on the average urban association index or “UAI” (based on levels of urban tolerance) of their species assemblages. To assess this metric, we used iNaturalist data to estimate a species-level UAI for 967 species across six broad taxonomic focal groups that occur in Southern California. We then applied these indices to spatiotemporally thinned species occurrence data in order to calculate the CUTI for a spatial grid covering the city of Los Angeles. The CUTI represents the degree that the terrestrial animal community is composed of species that are either tolerant or intolerant to urbanization. We then calculated a mean CUTI across all grid cells in Los Angeles, resulting in a single score for Metric 1.2b in the LA City Biodiversity Index. The methodology provided herein provides a framework for establishing repeat measures over time of urban tolerance within the city of Los Angeles and is applicable to other urban areas. Ultimately, these methods can help local managers and city officials across the region, state, or country understand and track the success (or failures) of local initiatives to support biodiversity and attract historically urban intolerant species to their cities. As urbanization poses a continued threat to biodiversity, particularly in biodiversity hotspots, the methods presented here will enable local governments to better manage and protect native biodiversity.

## METHODS

### Study Area

Our study was focused on Southern California, with an emphasis on the greater Los Angeles area, situated in the California Floristic Province, one of 36 biodiversity hotspots in the world (Myers et al. 2000; CEPF 2017). While the calculation of a formal CUTI was limited to areas within the city of Los Angeles, for the estimation of species-level UAI, our study area included all land within a 200 km buffer of the Los Angeles City boundary (approximate centroid: 34.031656, -118.241716), including the cities of Los Angeles, San Diego, Bakersfield, and Santa Barbara. We chose to focus on this broad geographic region because we were interested in creating a metric that could be measured repeatedly over time and would be robust to species that do not currently reside in our focal area of Los Angeles, but could colonize in the future. Furthermore, we treat urban tolerance here (as measured by the UAI) as a species-level trait, which is best estimated using occurrence data from a broader geographic region than just Los Angeles. As such, we aimed to include areas with a wide range of levels of urbanization, including multiple manifestations (i.e., cities) of urban land-use within the broader Southern California region.

### Urban Intensity

In order to estimate urban associations, we first had to define a continuous spatial layer of the urban intensity of our study region. To do this, we used Principal Component Analysis (PCA), which decomposes multivariate datasets into major axes of variation, to create a single composite index of urban intensity from multiple sources. Following Callaghan et al. (2020), this index included the Visible Infrared Imaging Radiometer Suite (VIIRS) nighttime lights data layer, but we added additional environmental variables related to urbanization as different taxa are likely to respond to different aspects of the urban environment. We initially tested a set of six data layers to depict urbanness across our study area, but we removed several layers (PM2.5, Average Annual Traffic Volume, and Population Density) due to collinearity and coarser resolution. Thus, our PCA index represents a composite of three layers: (1) light pollution from VIIRS Version 4 DMSP-OLS Nighttime Lights Time Series (https://eogdata.mines.edu/products/dmsp/#v4_dmsp_download) (Elvidge et al. 2013), (2) “Percent Impervious Surfaces” from National Land Cover Database (NLCD) 2016 Percent Developed Imperviousness (CONUS) (https://www.mrlc.gov/data/nlcd-2016-percent-developed-imperviousness-conus) (Dewitz 2019), and (3) “Noise Pollution” from National Park Service (NPS) Geospatial Sound Modeling 2013-2015 (https://irma.nps.gov/DataStore/Reference/Profile/2217356) (Mennitt et al. 2014). All spatial layers were reprojected and resampled, as needed, to a 0.25 x 0.25 mile grid prior to combination. The first axis of the PCA explained 86.5% of the variance in all three layers (**S1 Table**), indicating as expected that the three layers are all indicative of the same general process (i.e., “urbanness”) yet individually add unique information. Because PCA axis 1 (“PC1”) explained >70% of the variation, we retained it as our sole spatial index of urban intensity (**Fig. 1**).

**Figure 1.**
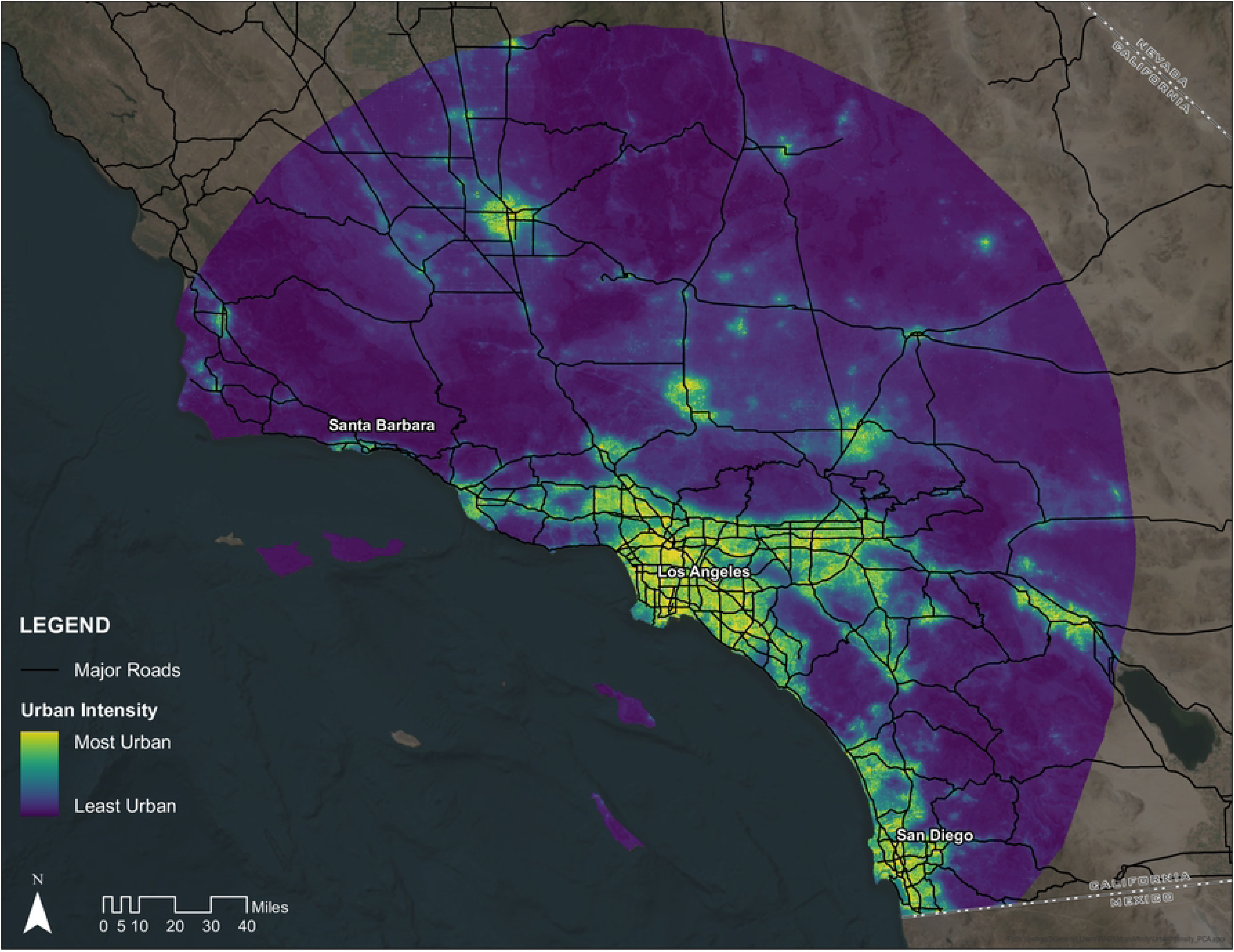
Urban intensity was measured across a broad study region in Southern California, defined by a 200 km radius circle centered on Los Angeles. Our metric of urban intensity was determined as the first PCA axis of three different variables. Warmer colors indicate higher levels of urban intensity. Solid lines detail major roadways within California.

### iNaturalist Records and Data Quality Filtering

We focused our analysis on select target taxonomic groups, which were picked a priori with expert input to represent 12 taxonomic groups that are generally well-detected and well-surveyed by community scientists on the iNaturalist platform. The 12 groups include: snails and slugs (Order: Stylommatophora); spiders (Order: Araneae); dragonflies and damselflies (Order: Odonata); grasshoppers, locusts, and crickets(Order: Orthoptera); leafhoppers (Family: *Cicadellidae*); lady beetles (Family: *Coccinellidae*); hoverflies (Family: *Syrphidae);* bees and wasps (Family: *Apidae* and *Vespidae*); butterflies and moths (Families: *Papilionidae, Pieridae, Lycaenidae, Nymphalidae, Sphingidae* and *Erebidae*); herpetofauna (Classes: Amphibia and Reptilia); mammals (Class: Mammalia); and birds (Class: Aves). We queried the iNaturalist API for occurrence data on 13 January 2022 using the ‘rinat’ package version 0.1.8 (Barve et al. 2022). We downloaded all iNaturalist records between 2011–2021 for the higher order taxa groups defined above, only limiting records to “research quality” georeferenced occurrences (i.e., those with a consensus taxonomic ID and location coordinates) bound within our broader study region (i.e., 200 km radius around Los Angeles). After downloading, we further filtered our data to remove species that had more than 60% of all observations records marked as “Geoprivacy = obscured” (a situation where iNaturalist provides spatial coordinates of sightings, but these coordinates are randomly offset by up to 22 km from the true location). Although we additionally filtered out all obscured spatial records for all species, we wholly excluded species meeting this arbitrary threshold as we believed that such a widespread degree of geoprivacy indicated species for which remaining iNaturalist data would not likely represent the species’ true distribution within the study area. Finally, we reclassified all records identified to the subspecies level to the species level following other similar work using related datasets (Feng and Che-Castaldo 2021; Neate-Clegg et al. 2023).

### Expert Review

After downloading iNaturalist records and applying our initial hard filters for data quality, we engaged the LASAN Biodiversity Expert Council, a regional group of engaged scientists and taxonomic specialists who advise the annual biodiversity report, to further assist in data curation and QA/QC. Experts in areas of specific taxonomic focus for the species included in this study were asked to evaluate occurrence data for all higher order taxonomic groupings using the following: 1) Is this species native to the study area; 2) Is the species terrestrial; 3) Is the natural history of this species so different from others in its taxonomic grouping, that records for this particular species should not be used as indicators of search effort for other similar species; and 4) Is there any other reason why we should exclude this species from this study. The third question refers to the issue that iNaturalist data are presence-only and do not, on their own, provide information on absence or non-detection. Increasingly, however, ecologists are using multi-taxa presence-only surveys to bin species into ‘detection groups’, whereby an observation of one species at a location provides inference on the non-detection of other species (Isaac et al. 2014; Outhwaite et al. 2020). This assumption of substitutability is justified as natural history observers are often searching broadly within taxonomic groups; for example, a birdwatcher’s positive record of one bird species says more about the non-detection of another bird species than it does about the non-detection of a butterfly. In the context of the present study, we did not require species’ occurrences to be perfect indices of non-detection for other species, but simply sought taxonomic groups where the presence of one species in that group would serve as a broad index of survey effort for all species in that group. Thus, we sought via expert review to exclude taxa that differed so much from the rest of their grouping (e.g., diurnal versus nocturnal; or identifiable via photography versus identifiable only via microscope) that they should not be treated as survey effort proxies. For the fourth question, some common reasons for excluding species based on expert review included species with extremely limited distributions that would otherwise be uninformative to urban tolerance (e.g., a species of plethodontid salamander limited to a single remaining population on Mt. Baldy, Los Angeles, USA), or misidentifications based on recent taxonomic splits.

Following data review by taxonomic specialists, we curated their responses to make sure that experts interpreted these questions similarly. We filtered observations based on these responses to exclude non-native species, species unlikely to be detected by typical observers, non-terrestrial (i.e. marine or freshwater) species, and species according to additional criteria as determined by the taxonomic group specialists.

### Controlling for Differences in Sampling Effort

We took a number of steps to control for differences in sampling evenness and effort in iNaturalist data. First, to address the inherent biases associated with sampling evenness, we performed a broad spatiotemporal thin (Boria et al. 2014, Steen et al. 2021). We thinned species-specific data to one observation per year within each 0.25 x 0.25 mile grid cell. This produced a database where every species is recorded as present or not detected in each grid cell and for each year between 2011–2021 (i.e. the number of yearly detections out of 11 years). Second, to address the additional bias associated with varied sampling effort, we defined site-specific sampling effort for each of the twelve focal taxonomic groups. We did this because observers may not equally record observations for all taxonomic groups, (e.g., an observer may not observe spiders while recording birds).Taxonomic group-specific effort grids were calculated based on the observation effort per year for the corresponding focal taxonomic groups, summing within grid cells the number of years with at least one observation in a year of a species from within a taxonomic group. Thus, taxonomic group-specific effort layers indicate the number of years (0–11) in which observers recorded at least one record of a target group, which serves as the maximum potential number of thinned presences for any given species of that group. As such, the thinned species presence layer and the matching effort layer represent a spatially-varying binomial response, where the number of binomial trials is the effort in a grid cell and the number of binomial successes is the thinned number of species’ presences.

### Measuring Species-level Relationship to Urban Intensity Layer in Study Area

Species-level indices of urban association were calculated based on thinned occurrence data and taxon-matched effort data across the entire Southern California study region. We calculated a UAI for each species that had at least 25 thinned annual occurrences across our study region. Specifically, using the ‘stats’ package (R Core Team 2022), for each species we modeled the number of thinned occurrences, given effort per cell, as a binomial process that varied as a function of a single covariate: the urban intensity (PC1) of the grid cell. This model allowed us to estimate the number of thinned species occurrences as a binomial variable, where the number of successes (i.e., ‘occurrences’) was capped by the number of years with non-zero effort for the taxonomic group in each cell. In this way, our model accounted for taxon-specific sampling effort over time in each grid cell. The resulting slope, which indicates the relationship between species’ occurrence and urban intensity, was stored as the UAI for a given species. Positive slopes indicate urban tolerance, negative slopes suggest urban intolerance, and slopes of zero indicate no relationship of occurrence to urban intensity.

### Calculating a Community Urban Tolerance Index (CUTI)

After calculating a UAI for each species, we quantified a CUTI for each grid cell in Los Angeles by taxonomic group, as well as a composite score for the entire city (i.e., metric 1.2b for the city of Los Angeles). To calculate a taxonomic group-specific CUTI or each grid cell in Los Angeles, we matched species’ UAIs to species occurrences in individual grid cells and calculated a raw CUTI score per cell by taking the mean UAI score of all species within any given cell, weighted by the thinned temporal occurrence of each species (i.e., a value of 1–11 for the number of years that the species occurred in that cell). This resulted in 12 grids (one for each taxonomic group) with a group-specific CUTI score for every cell in the city in which the group was detected. To interpret these results at a broader taxonomic scale, we also calculated the mean CUTI across all pixels for each of the 12 taxonomic groups. Finally, to calculate a composite CUTI across all taxonomic groups, we averaged the 12 taxonomic group grids and city-wide scores. In all cases, the raw CUTI scores were binned into a 5-point scale as follows: -Infinity to -0.5 = 5; -0.5 to -0.25 = 4; -0.25 to 0 = 3; 0 to 0.25 = 2; 0.25 to 0.5 = 1; and 0.5 to Infinity = 0. On this scale, a cell with a CUTI index of 4 or 5 suggests that species in aggregate are more natural-area associated, while a cell with an index of 0 or 1 suggests that species are more urban tolerant. To test for an association between UIA and urban intensity (with the hypothesis that areas of higher urban intensity have lower CUTI scores), we used an ANOVA and Tukey HSD test in the ‘stats’ package (R Core Team 2022) with urban intensity as the response variable and categorical binned CUTI scores for each grid cell as an independent variable.

## RESULTS

Our iNaturalist query yielded a total of 958,624 observations from 127,553 observers (**Table 1**). After filtering these observations, we retained 567,996 observations from 71,120 observers. Our filtered query included a total of 967 unique native species found within the study area, of which 563 occurred at least once within the city of Los Angeles. We were able to calculate UAI for 510 species in our dataset, of which 408 occurred at least once within the city of Los Angeles. The species assessed were on average negatively associated with our measure of urbanization, although there was variation across species (cross-species mean = - 0.21, range = -2.93 to 0.62; Fig. 1, Fig. 2**, & S2 Table**). UAI varied between the 12 taxonomic groupings (**Table 2**), with snails and slugs having the highest (i.e. more urban tolerant) score (group mean = 0.24, range = -0.096 to 0.62), and butterflies and moths having the lowest (i.e. more urban intolerant) UAI (group mean = -0.40, range = -2.93 to 0.46). The most urban associated species in our study was the slipper snail (*Cochlicopa lubrica*) (UAI = 0.62), and the least urban associated species was the greenish blue butterfly (*Icaricia saepiolus*) (UAI = -2.93).

**Figure 2.**
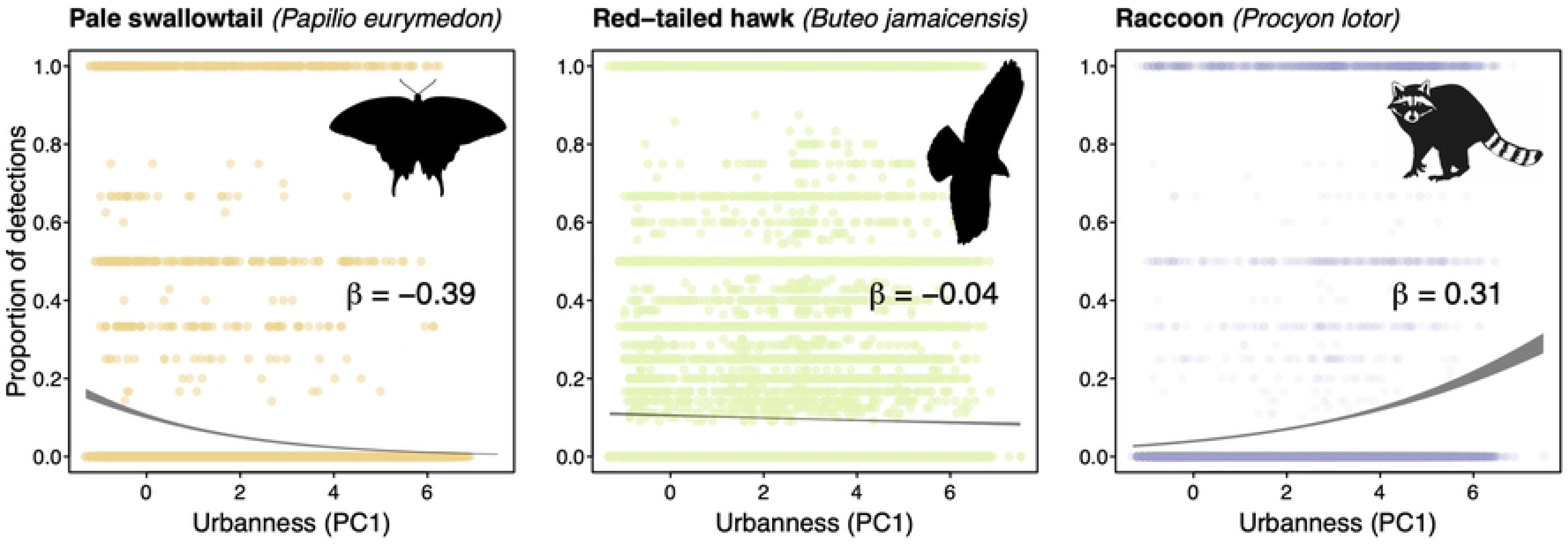
Species vary widely in their response of occurrence to urbanization, as exemplified by three different taxa showing urban intolerance (left), urban ambivalence (center), and urban tolerance (right). Scatterplots show the proportion of detections (out of a maximum of 11 years) for each species across each grid cell in the broader Southern California study region (Fig. 1). Trend lines show the 95% confidence interval surrounding a binomial regression of detection frequency as a function of urbanness. Species’ UAI scores (*β*) are the logit-linear slope of the trend line.

**Table 1.** Counts of observations and observers between unfiltered dataset downloaded from iNat API and iNaturalist data subject to exclusion by expert review (see methods for criteria) and hard filters.

|  |  |  | Unfiltered |  | Filtered |  |  |
| --- | --- | --- | --- | --- | --- | --- | --- |
| Higher Order Grouping | Taxon | iNat Taxa ID | # Observations | # Observers | # Observations | # Observers | # Species |
| Snails and Slugs | *Stylommatophora* | 47485 | 22,539 | 5,507 | 919 | 213 | 5 |
| Spiders | *Araneae* | 47118 | 28,148 | 8,897 | 16,029 | 5,778 | 144 |
| Dragonflies and Damselflies | *Odonata* | 47792 | 16,391 | 3,431 | 16,366 | 3,426 | 73 |
| Grasshoppers, Locusts, and Crickets | *Orthoptera* | 47651 | 17,267 | 5,670 | 13,816 | 4,483 | 140 |
| Leaf Hoppers | *Cicadellidae* | 53237 | 2,585 | 849 | 796 | 236 | 41 |
| Lady Beetles | *Coccinellidae* | 48486 | 18,030 | 5,090 | 9,246 | 2,677 | 37 |
| Hoverflies | *Syrphidae* | 49995 | 10,654 | 2,200 | 8,212 | 1,832 | 40 |
| Bees and Wasps | *Apidae* | 47221 | 34,748 | 11,105 | 11,499 | 3,166 | 68 |
|  | *Vespidae* | 52747 | 5,113 | 1,826 |  |  |  |
| Butterflies and Moths | *Erebidae* | 121850 | 3,939 | 1,714 | 28,480 | 6,973 | 68 |
|  | *Lycaenidae* | 47923 | 15,199 | 2,479 |  |  |  |
|  | *Nymphalidae* | 47922 | 38,786 | 10,023 |  |  |  |
|  | *Papilionidae* | 47223 | 8,514 | 3,963 |  |  |  |
|  | *Pieridae* | 48508 | 9,528 | 2,504 |  |  |  |
|  | *Sphingidae* | 47213 | 6,988 | 3,967 |  |  |  |
| Herpetofauna | *Amphibia* | 20978 | 19,892 | 5,229 | 72,312 | 13,992 | 42 |
|  | *Reptilia* | 26036 | 70,637 | 13,601 |  |  |  |
| Mammals | *Mammalia* | 40151 | 56,914 | 11,763 | 34,107 | 7,514 | 49 |
| Birds | *Aves* | 3 | 572,752 | 27,735 | 356,214 | 20,830 | 260 |
| **TOTAL** |  |  | 843,010 | 95,909 | 502,612 | 52,475 | 487 |

**Table 2.** Average urban association index (UAI) scores for each of the 12 higher order taxonomic groupings. Species-level UAI scores ranged from -2.9 to 0.62, with more negative numbers indicating more urban intolerant species and more positive scores indicating more urban tolerant species.

| Higher Order Grouping | Average UAI Score |
| --- | --- |
| Snails and Slugs | 0.24 |
| Spiders | -0.14 |
| Dragonflies and Damselflies | -0.20 |
| Grasshoppers, Locusts, and Crickets | -0.37 |
| Leaf Hoppers | -0.08 |
| Lady Beetles | -0.08 |
| Hoverflies | -0.11 |
| Bees and Wasps | -0.16 |
| Butterflies and Moths | -0.40 |
| Herpetofauna | -0.34 |
| Mammals | -0.39 |
| Birds | -0.15 |

We assessed 8,348 0.25 x 0.25 mile grid cells across the city of Los Angeles for their CUTI, weighted by the temporally thinned occurrence of each species. A total of 2,010 grid cells did not have records for any target taxa after filtering and were not included in our calculation of summary scores. Averaging across all higher order taxonomic groupings, the city had an average CUTI of 2.01 (raw, unbinned score = 0.129; Fig. 3). Average CUTI varied between higher order taxonomic groupings (**Table 3)**, with snails and orthoptera demonstrating the highest average IUM (snails: 1.40 binned, 0.30 raw; orthoptera: 1.54 binned, 0.17 raw), and odonates and mammals showing the lowest average CUTI (odonates: 2.25 binned, 0.06 raw; mammals: 2.16 binned, 0.07 raw) (Fig. 4). There was a significant relationship between urbanization values and the cross-taxa average CUTI of those cells, with areas of higher urban intensity holding taxa that, on the whole, were more urban tolerant (i.e, have higher UAI values) (ANOVA, p < 0.001; **S1 Figure**). This general relationship held true for every individual taxonomic group (ANOVA, p < 0.001), except for snails (ANOVA, p = 0.99). There was a weak positive relationship between

**Figure 3.**
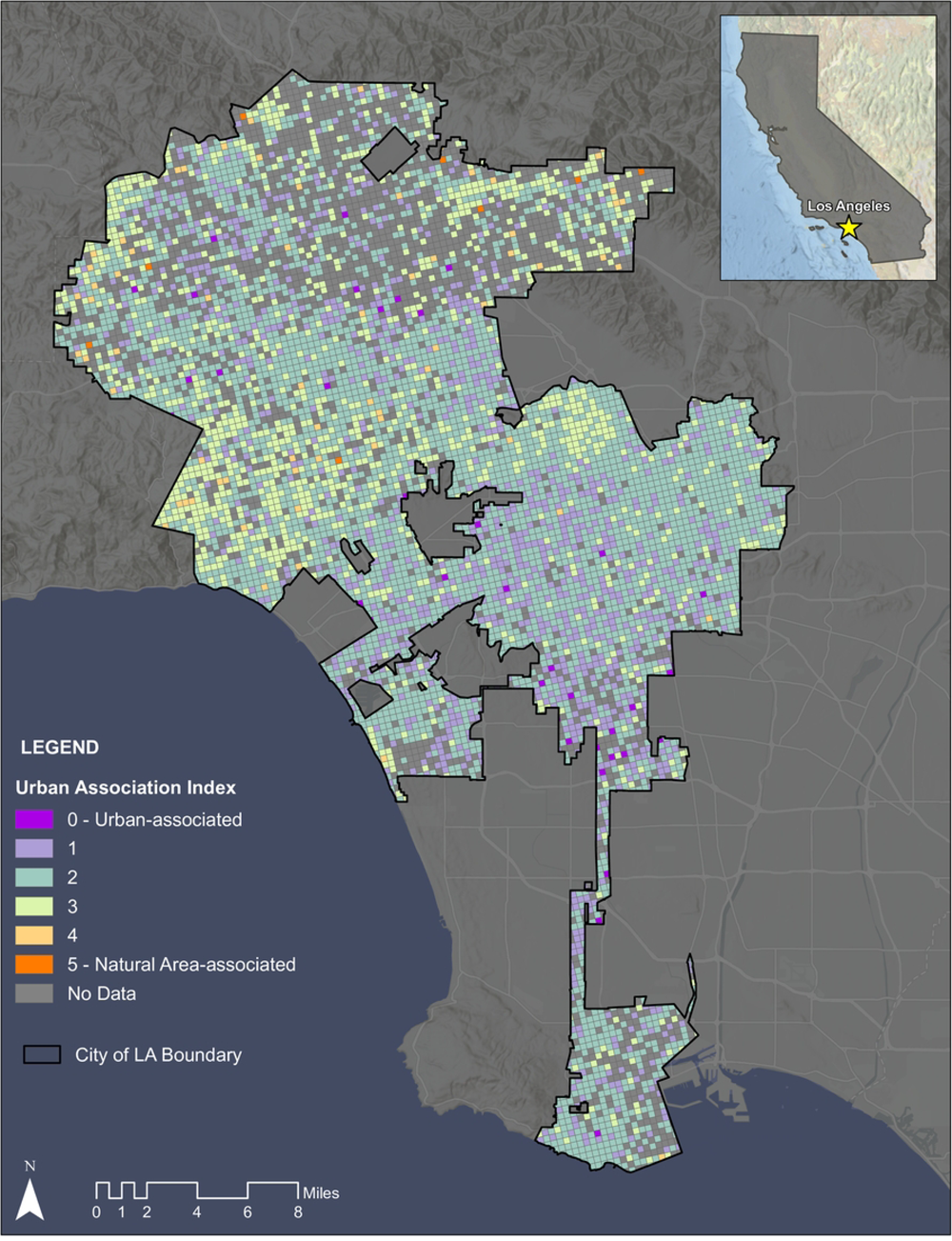
Map of the City of Los Angeles with overlaid mean composite urban association scores on the 0.25 x 0.25 mile scale. Warmer colors indicate more natural area tolerant species, whereas cooler colors indicate more urban tolerant species. Areas within Los Angeles city boundaries with insufficient data to calculate the score are colored gray.

**Figure 4.**
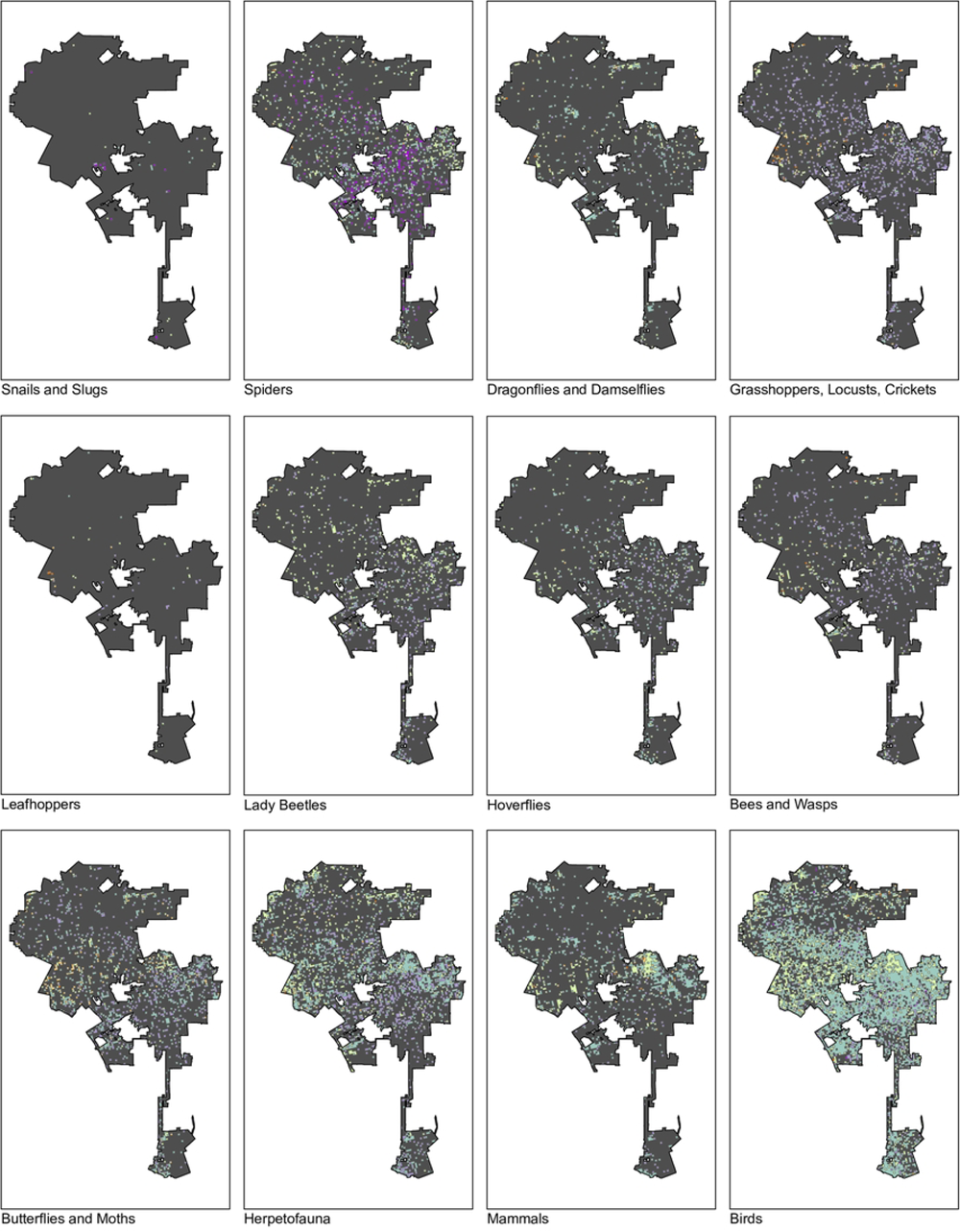
Map of the urban association scores separated by higher order taxonomic groupings. Levels correspond to Figure 2, where warmer colors indicate more urban intolerant species, cooler colors indicate more urban tolerant species, and gray indicates cells with insufficient data to calculate a score.

**Table 3.**
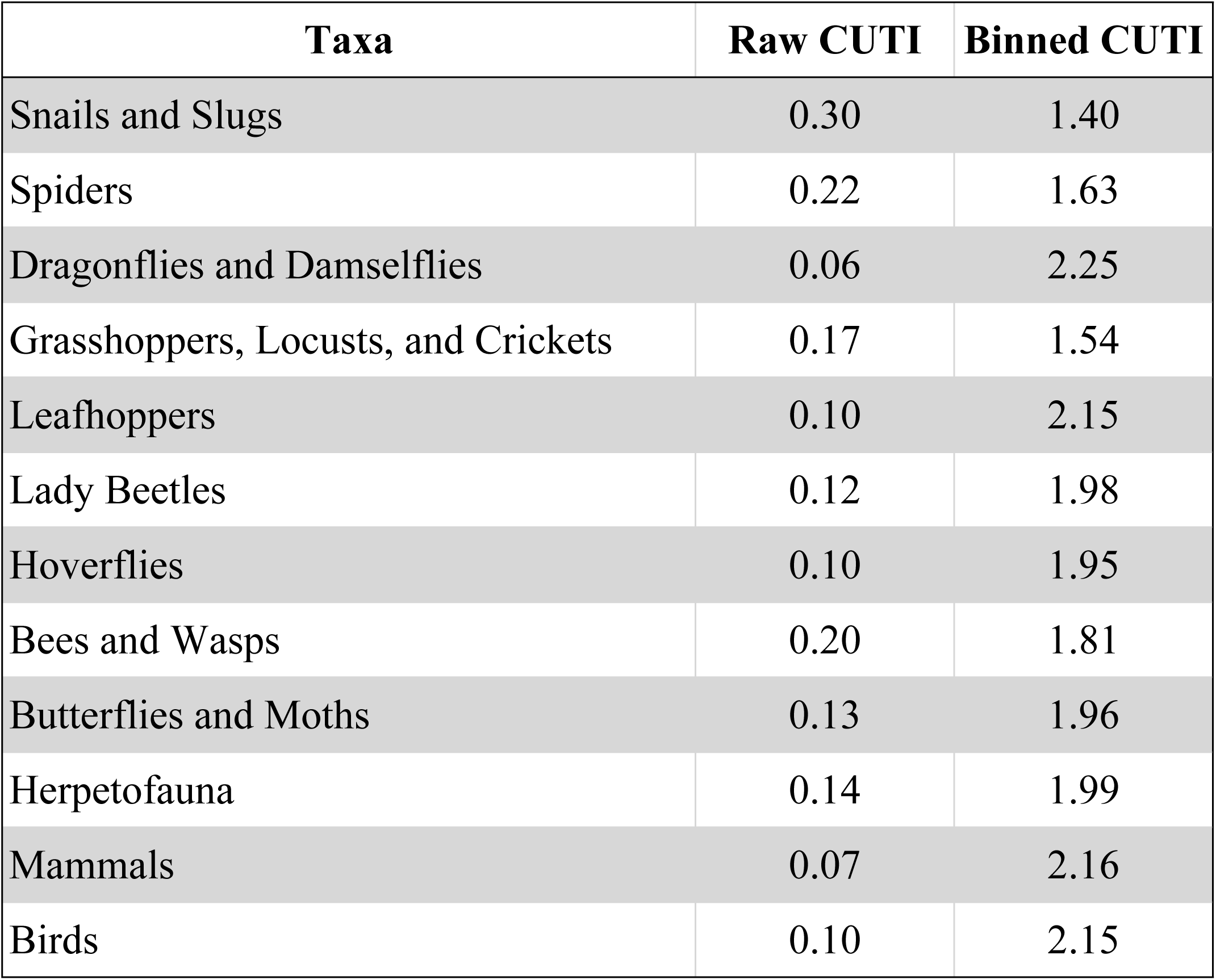
Community urban tolerance index (CUTI) scores for each of the 12 higher order taxonomic groupings across the study area. Raw values are the average of weighted average of species-level UAI scores, while binned values rescale to a 5-point index, where a CUTI index of 4–5 suggests that species in aggregate are more natural-area associated, while a an index of 0–1 suggests that species are more urban tolerant.

| Taxa | Raw CUTI | Binned CUTI |
| --- | --- | --- |
| Snails and Slugs | 0.30 | 1.40 |
| Spiders | 0.22 | 1.63 |
| Dragonflies and Damselflies | 0.06 | 2.25 |
| Grasshoppers, Locusts, and Crickets | 0.17 | 1.54 |
| Leafhoppers | 0.10 | 2.15 |
| Lady Beetles | 0.12 | 1.98 |
| Hoverflies | 0.10 | 1.95 |
| Bees and Wasps | 0.20 | 1.81 |
| Butterflies and Moths | 0.13 | 1.96 |
| Herpetofauna | 0.14 | 1.99 |
| Mammals | 0.07 | 2.16 |
| Birds | 0.10 | 2.15 |

## DISCUSSION

Using over 567,000 publically available community science records from iNaturalist, we present the first comprehensive species-level evaluation of urban tolerance for Southern California taxa. For the City of Los Angeles, we found that on average, native species within the city were negatively associated with our measure of urbanness, and that this pattern was conserved across the higher order taxonomic groupings with the exception of snails and slugs, which were positively associated with our measure of urbanness. We then averaged this species-level response to our measure of urban intensity on a ¼ mile grid across the City of Los Angeles to understand the geographic distribution of urban intolerant species. Ultimately, this metric of native species urban tolerance (CUTI) can be reassessed regularly as a means of evaluating change in urban tolerance over time and following specific biodiversity improvement measures in the city.

The most urban tolerant species in our study was the slipper snail (*C. lubrica)*. This species is known to be widespread and euryhygric (i.e. able to withstand a broad range of moisture conditions), and it is possible that urbanized areas such as Los Angeles provide year-round access to a variety of moisture regimes through the addition of ornamental landscapes and lawns (Barker 1999; Georgiev 2008; Čejka and Hamerlík 2009). Other studies have found high native snail abundance in areas of high urban intensity in Tennessee (Hodges and McKinney 2018). Given that there were only five species of snails and slugs that remained in our dataset post-filtering, this pattern could largely be due to a small sample size, and perhaps more purposeful sampling is required to truly ascertain the affinity of this group for urban environments. All other higher order taxonomic groups had higher presence in less urbanized regions within the city. In Los Angeles, research has demonstrated that the presence of many taxonomic groups is negatively affected by increased levels of urbanization. For example, coyotes and bobcats in mixed urban/natural areas have home ranges that primarily utilize natural areas (Riley et al. 2003), and some regionally common amphibian species are markedly absent from streams within urbanized areas of the city (Riley et al. 2005).

The most urban intolerant species in our study was the greenish blue butterfly (*Icaricia saepiolus)*. Compared to other North American butterfly families, *Lycaenidae* is overrepresented in terms of number of species proposed for listing (Cushman and Murphy 1993) largely due to their host plant specificity, primarily for plants in the genera *Lupinus* and *Eriogonum,* and the fact that these plants are adapted to disturbance regimes that are infrequent in the urban context (Cushman and Murphy 1993; MacDonald et al. 2012; Longcore and Osborne 2015). Conservation of many of the special status butterfly species including several Lycaenids therefore relies on maintaining and expanding critical segments of habitat that contain host plant species within urban settings and maintaining habitat fragments of varying sizes through deliberate disturbance as a management tool (Longcore and Osborn 2015). For example, the Palos Verdes blue butterfly (*Glaucopsyche lygdamus palosverdesensis*) is an endangered subspecies of Lycaenid butterfly in Los Angeles County, and findings from the US Fish and Wildlife Service demonstrate that the species appears to be establishing in reintroduction sites due primarily to efforts to remediate historical habitat through mechanical disturbance, non-native plant removal, and planting of successional host plants (Anderson 2014). Based on the findings from this work, the City of Los Angeles could use existing observations to identify target areas for conservation. In areas where observations overlap with property owned and managed by the City of Los Angeles, the city can focus restoration efforts on increasing host plant abundance through direct plantings of early successional plants and targeted mechanical disturbance to create conditions necessary for early successional habitat needed by the butterflies. Following restoration activities like these, city managers can evaluate success through a reevaluation of this metric on an annual basis.

While this study includes over 500 native species observed within the study area, approximately 59% of the total community science records were excluded from the analyses because they were considered undetectable to the general public (e.g. small insects, nocturnal mammals, etc.), the data was not at the “research grade” level, or the records for a given species were geoobscured due to species status or user preference. Other studies have suggested similar data quality issues and biases in iNaturalist records (Hochmair et al. 2020; Mesaglio and Callaghan 2021; Chesshire et al. 2023). In order to circumvent data quality issues, this study relied on expert review to identify higher order taxonomic groupings that could be reliably identified by the general public, but in doing so, may have increased ascertainment bias and decreased the overall scope of the data. In an effort to reduce the amount of data lost, future assessments of this metric may benefit from developing a relationship with iNaturalist in order to obtain user-obscured data en masse, which is not currently possible without requesting thousands of individual records from each iNaturalist user.

While we present several limitations to the available crowd-sourced species presence dataset within our sampling area, these data limitations also provide targets for local environmental managers to improve these datasets and therefore biodiversity monitoring in their regions. Based solely on the number of assessed grid cells across the City of Los Angeles in this study, it is clear that there needs to be substantial effort placed on bolstering community science projects that focus on underrepresented taxonomic groups (e.g. snails and slugs and leafhoppers). Findings from other community science projects indicate that local city residents are underrepresented contributors to community science datasets (Dimson and Gillepsie 2023), yet for community science based biodiversity monitoring to be successful, it must be built from a bottom-up approach that includes both participatory and contributory opportunities for the communities where biodiversity monitoring is to take place (Pocock et al. 2018). While some efforts in Los Angeles to involve residents in taxonomically focused community science projects have led to increased knowledge of urban biodiversity for these taxa groups (e.g. the BioSCAN project, see https://nhm.org/community-science-nhm/bioscan) (Brown and Hartop 2017), these projects are limited in geographic scope, are often short-term assessments, and the primary role of residents is that of data collection, which may not lead to sustained participation by community members in the future (West and Pateman 2016). Moreover, efforts to educate the public on specific indicator species is underway in Los Angeles (LASAN 2022b), and these efforts could be directed at underrepresented taxonomic groups highlighted in this study. It has been demonstrated that community science datasets can match if not surpass traditional biodiversity assessment methods in data quantity, and do so in a fraction of the time (Kittelberger et al. 2021; Shirey et al. 2021). Therefore, developing long-term and mutually beneficial partnerships with local communities to assess urban biodiversity should be a primary focus of city managers who plan to use large unstructured community science datasets to measure the efficacy of city-wide biodiversity measures.

Herein we present a broad taxonomic assessment of the urban tolerance of native animal species for the City of Los Angeles. We found that there are clear differences in species level responses to our measures of urban intensity and that native species within Southern California are largely urban intolerant. This is even more true within the City of Los Angeles. This study provides a baseline assessment of the degree of presence of urban intolerant species within the City of Los Angeles in a ten year study period. Repeated assessment of this metric will allow stakeholders such as the City of Los Angeles to monitor success of its stated goal of no-net biodiversity loss by 2035. An important metric within the City’s Biodiversity Index is the ability of this urban system to attract and maintain healthy populations of urban intolerant species.

## ACKNOWLEDGEMENTS

We would like to acknowledge Dan Cooper, Miguel Ordenana, and Jessica West for providing us access to iNaturalist project data. The study authors would like to specifically thank Gary Bucciarelli, Rachel Chock, and Jann Vendetti for their expert review of inaturalist data of herpetofauna, mammals, and slugs and snails, respectively. Finally, the study authors would like to acknowledge the LA City Biodiversity Expert Council, and particularly the members who attended the workshop for Metric 1.2B.

## AUTHOR CONTRIBUTIONS

J.N.C., M.B., R.G.F., M.L., A.L., and A.Y.P. designed the project and oversaw early iterations of the project. M.W.T. designed and implemented all statistical analysis and modeling for the project. G.A.M. performed expert review for all invertebrate taxa. K.R. designed all study maps. J.N.C. wrote the manuscript with assistance from M.B., R.G.F, M. L, A.L., G.A.M., A.Y.P., and M.W.T. All authors reviewed and commented on the manuscript prior to submission.

## SUPPLEMENTAL MATERIAL

Supplemental material including all scripts used to generate results and supplemental figures and tables are uploaded and embargoed on Zenodo. These materials will be made available upon publication.

---

**Supplementary Table 1.** Table detailing the loadings and variance explained for the composite urban intensity layer separated by contributing layers and PC axes.

**Supplementary Table 2.** Table of all species considered in this paper and their associated urban affinity scores.

**Supplementary Figure 1.** Regression plot of urban intensity (first PCA axis) and CUTI scores for all 510 species.

## Notes

### Competing Interest Statement

The authors have declared no competing interest.

## REFERENCES

1. Ancillotto L, Santini L, Ranc N, Maiorano L, Russo D. Extraordinary range expansion in a common bat: the potential roles of climate change and urbanization. The Science of Nature. 2016 Apr;103(3–4):15.

2. Anderson A. Palos Verdes blue butterfly (*Glaucopsyche lygdamus palosverdesensis*) 5-Year Review. United States Fish and Wildlife Service; 2014. Available from: https://ecosphere-documents-production-public.s3.amazonaws.com/sams/public_docs/species_nonpublish/2149.pdf

3. Barker GM. Naturalized terrestrial Stylommatophora: Mollusca: Gastropoda. Lincoln: Manaaki Whenua Press; 1999. 253 p. (Fauna of New Zealand).

4. Barve V, Hart E, Guillou S. rinat: Access’ iNaturalist’data through APIs. R package version 01. 2022;8.

5. Batáry P, Kurucz K, Suarez-Rubio M, Chamberlain DE. Non-linearities in bird responses across urbanization gradients: A meta-analysis. Global Change Biology. 2018 Mar;24(3):1046– 54.

6. Bayraktarov E, Ehmke G, O’Connor J, Burns EL, Nguyen HA, McRae L, et al. Do Big Unstructured Biodiversity Data Mean More Knowledge? Frontiers in Ecology and Evolution. 2019 Jan 24;6:239.

7. Beninde J, Veith M, Hochkirch A. Biodiversity in cities needs space: a meta-analysis of factors determining intra-urban biodiversity variation. Ecol Lett. 2015 Jun;18(6):581–92.

8. Blair R. The Effects of Urban Sprawl on Birds at Multiple Levels of Biological Organization. Ecology and Society. 2004;9(5):art2.

9. Blair RB, Johnson EM. Suburban habitats and their role for birds in the urban–rural habitat network: points of local invasion and extinction? Landscape Ecology. 2008 Dec;23(10):1157–69.

10. Boria RA, Olson LE, Goodman SM, Anderson RP. Spatial filtering to reduce sampling bias can improve the performance of ecological niche models. Ecological Modelling. 2014 Mar;275:73–7.

11. Brook B, Sodhi N, Bradshaw C. Synergies among extinction drivers under global change. Trends in Ecology & Evolution. 2008 Aug;23(8):453–60.

12. Brown BV, Hartop EA. Big data from tiny flies: patterns revealed from over 42,000 phorid flies (Insecta: Diptera: Phoridae) collected over one year in Los Angeles, California, USA. Urban Ecosystems. 2017 Jun;20(3):521–34.

13. Callaghan CT, Ozeroff I, Hitchcock C, Chandler M. Capitalizing on opportunistic citizen science data to monitor urban biodiversity: A multi-taxa framework. Biological Conservation. 2020 Nov;251:108753.

14. Ceballos G, Ehrlich PR, Barnosky AD, García A, Pringle RM, Palmer TM. Accelerated modern human–induced species losses: Entering the sixth mass extinction. Science Advances. 2015 Jun 5;1(5):e1400253.

15. Čejka T, Hamerlík L. Land snails as indicators of soil humidity in Danubian woodland (SW Slovakia). Polish Journal of Ecology. 2009;57(4):741–7.

16. Chesshire PR, Fischer EE, Dowdy NJ, Griswold TL, Hughes AC, Orr MC, Ascher JS, Guzman LM, Hung KL, Cobb NS, McCabe LM. Completeness analysis for over 3000 United States bee species identifies persistent data gap. Ecography. 2023 Feb 6:e06584.

17. Cushman J, Murphy D. Conservation of North American lycaenids – an overview. In: Conservation Biology of Lycaenidae(Butterflies). Gland, Switzerland: IUCN; 1993:37-44.

18. (CEPF) Critical Ecosystem Partnership Fund. The North American Coastal Plain Recognized as the World’s 36th Biodiversity Hotspot. 2017. Available from: https://www.cepf.net/node/5472\

19. de Toledo MCB, Donatelli RJ, Batista GT. Relation between green spaces and bird community structure in an urban area in Southeast Brazil. Urban Ecosystems. 2012 Mar;15(1):111– 31.

20. Dewitz J. National land cover database (NLCD) 2016 products: US Geological Survey data release. 2019

21. Di Cecco GJ, Barve V, Belitz MW, Stucky BJ, Guralnick RP, Hurlbert AH. Observing the Observers: How Participants Contribute Data to iNaturalist and Implications for Biodiversity Science. BioScience. 2021 Nov 2;71(11):1179–88.

22. Dimson M, Gillespie TW. Who, where, when: Observer behavior influences spatial and temporal patterns of iNaturalist participation. Applied Geography. 2023 Apr;153:102916.

23. Elvidge CD, Baugh KE, Zhizhin M, Hsu FC. Why VIIRS data are superior to DMSP for mapping nighttime lights. APAN Proceedings. 2013 Jun 10;35(0):62.

24. Faeth SH, Bang C, Saari S. Urban biodiversity: patterns and mechanisms: Urban biodiversity. Annals of the New York Academy of Sciences. 2011 Mar;1223(1):69–81.

25. Feng MLE, Che-Castaldo J. Comparing the reliability of relative bird abundance indices from standardized surveys and community science data at finer resolutions. Silva D de P, editor. PLoS ONE. 2021 Sep 10;16(9):e0257226.

26. Fenoglio MS, Calviño A, González E, Salvo A, Videla M. Urbanization drivers and underlying mechanisms of terrestrial insect diversity loss in cities. Ecological Entomology. 2021 Aug;46(4):757–71.

27. Fenoglio MS, Rossetti MR, Videla M. Negative effects of urbanization on terrestrial arthropod communities: A meta-analysis. Global Ecology and Biogeography. 2020 Aug;29(8):1412–29.

28. Fontana S, Sattler T, Bontadina F, Moretti M. How to manage the urban green to improve bird diversity and community structure. Landscape and Urban Planning. 2011 Jun;101(3):278–85.

29. Gandomi A, Haider M. Beyond the hype: Big data concepts, methods, and analytics. International Journal of Information Management. 2015 Apr;35(2):137–44.

30. Georgiev D. Habitat distribution of the land snails in one village area of the upper Thracian valley (Bulgaria). Anniversary Scientific Conference of Ecology, Proceedings. 2008:147–151.

31. Hochmair HH, Scheffrahn RH, Basille M, Boone M. Evaluating the data quality of iNaturalist termite records. Barden P, editor. PLoS ONE. 2020 May 4;15(5):e0226534.

32. Hodges MN, McKinney ML. Urbanization impacts on land snail community composition. Urban Ecosystems. 2018 Aug;21(4):721–35.

33. Husté A, Boulinier T. Determinants Of Local Extinction And Turnover Rates In Urban Bird Communities. Ecological Applications. 2007 Jan;17(1):168–80.

34. IPBES. Global assessment report on biodiversity and ecosystem services of the Intergovernmental Science-Policy Platform on Biodiversity and Ecosystem Services. 2019.

35. Isaac NJB, Strien AJ, August TA, Zeeuw MP, Roy DB. Statistics for citizen science: extracting signals of change from noisy ecological data. Anderson B, editor. Methods in Ecology and Evolution. 2014 Oct;5(10):1052–60.

36. Kamp J, Oppel S, Heldbjerg H, Nyegaard T, Donald PF. Unstructured citizen science data fail to detect long-term population declines of common birds in Denmark. Diversity and Distributions. 2016 Oct;22(10):1024–35.

37. Kittelberger KD, Hendrix SV, Şekercioğlu ÇH. The Value of Citizen Science in Increasing Our Knowledge of Under-Sampled Biodiversity: An Overview of Public Documentation of Auchenorrhyncha and the Hoppers of North Carolina. Frontiers in Environmental Science. 2021 Aug 27;9:710396.

38. (LASAN 2018) 2018 Biodiversity Report. Los Angeles Sanitation and Environment. 2018. Available from: https://www.lacitysan.org/cs/groups/public/documents/document/y250/mdi0/~edisp/cnt024743.pdf

39. (LASAN 2020) 2020 Biodiversity Report. Los Angeles Sanitation and Environment. 2020. Available from: https://www.lacitysan.org/cs/groups/public/documents/document/y250/mduy/~edisp/cnt052553.pdf

40. (LASAN 2022a) LA Biodiversity Index Baseline Report. Los Angeles Sanitation and Environment. 2022a. Available from: https://www.lacitysan.org/san/sandocview?docname=cnt076756

41. (LASAN 2022b) Biodiversity Indicator Species: A Guide to the City of Los Angeles’ Charismatic Umbrella Species. Los Angeles Sanitation and Environment. 2022b. Available from: https://www.lacitysan.org/cs/groups/public/documents/document/y250/mdc1/~edisp/cnt075161.pdf

42. Lerman SB, Warren PS. The conservation value of residential yards: linking birds and people. Ecological Applications. 2011 Jun;21(4):1327–39.

43. Li E, Parker SS, Pauly GB, Randall JM, Brown BV, Cohen BS. An Urban Biodiversity Assessment Framework That Combines an Urban Habitat Classification Scheme and Citizen Science Data. Frontiers in Ecology and Evolution. 2019 Jul 17;7:277.

44. Longcore T, Osborne KH. Butterflies Are Not Grizzly Bears: Lepidoptera Conservation in Practice. In: Daniels JC, editor. Butterfly Conservation in North America. Dordrecht: Springer Netherlands; 2015:161–92.

45. MacDonald B, Longcore T, Weiss S. Status and variability of mission blue butterfly populations at Milagra Ridge, Marin Headlands, and Oakwood Valley. The Urban Wildlands Group, Los Angeles. 2012 May 20:1–69.

46. Magle SB, Hunt VM, Vernon M, Crooks KR. Urban wildlife research: Past, present, and future. Biological Conservation. 2012 Oct;155:23–32.

47. McCallum ML. Vertebrate biodiversity losses point to a sixth mass extinction. Biodiversity and Conservation. 2015 Sep;24(10):2497–519.

48. McDonnell MJ, Pickett STA. Ecosystem Structure and Function along Urban-Rural Gradients: An Unexploited Opportunity for Ecology. Ecology. 1990 Aug;71(4):1232–7.

49. Mennitt D, Sherrill K, Fristrup K. A geospatial model of ambient sound pressure levels in the contiguous United States. The Journal of the Acoustical Society of America. 2014 May 1;135(5):2746–64.

50. Mesaglio T, Callaghan CT. An overview of the history, current contributions and future outlook of iNaturalist in Australia. Wildlife Research. 2021 Mar 19;48(4):289–303.

51. Munstermann MJ, Heim NA, McCauley DJ, Payne JL, Upham NS, Wang SC, et al. A global ecological signal of extinction risk in terrestrial vertebrates. Conservation Biology. 2022 Jun;36(3).

52. Myers N, Mittermeier RA, Mittermeier CG, da Fonseca GAB, Kent J. Biodiversity hotspots for conservation priorities. Nature. 2000 Feb;403(6772):853–8.

53. Neate-Clegg MH, Tonelli BA, Youngflesh C, Wu JX, Montgomery GA, Şekercioğlu ÇH, Tingley MW. Traits shaping urban tolerance in birds differ around the world. Current Biology. 2023 May;33(9):1677–1688.e6.

54. Outhwaite CL, Gregory RD, Chandler RE, Collen B, Isaac NJB. Complex long-term biodiversity change among invertebrates, bryophytes and lichens. Nature Ecology & Evolution. 2020 Feb 17;4(3):384–92.

55. Paker Y, Yom-Tov Y, Alon-Mozes T, Barnea A. The effect of plant richness and urban garden structure on bird species richness, diversity and community structure. Landscape and Urban Planning. 2014 Feb;122:186–95.

56. Parmesan C, Yohe G. A globally coherent fingerprint of climate change impacts across natural systems. Nature. 2003 Jan;421(6918):37–42.

57. Pocock MJO, Chandler M, Bonney R, Thornhill I, Albin A, August T, et al. A Vision for Global Biodiversity Monitoring With Citizen Science. In: Advances in Ecological Research. Elsevier; 2018:169–223.

58. Pocock MJ, Chandler M, Bonney R, Thornhill I, Albin A, August T, Bachman S, Brown PM, Cunha DG, Grez A, Jackson C. A Vision for Global Biodiversity Monitoring With Citizen Science. In: Bohan DA, Dumbrell AJ, Woodward G, Jackson M, editors. Next Generation Biomonitoring: Part 2. Academic Press; 2018. p. 169–223. (Advances in Ecological Research; vol. 59).

59. R Core Team. R: A Language and Environment for Statistical Computing. Vienna, Austria: R Foundation for Statistical Computing; 2022. Available from: https://www.R-project.org/

60. Rapacciuolo G, Young A, Johnson R. Deriving indicators of biodiversity change from unstructured community-contributed data. Oikos. 2021 Aug;130(8):1225–39.

61. Rega-Brodsky CC, Aronson MFJ, Piana MR, Carpenter ES, Hahs AK, Herrera-Montes A, et al. Urban biodiversity: State of the science and future directions. Urban Ecosystems. 2022 Aug;25(4):1083–96.

62. Riley SPD, Busteed GT, Kats LB, Vandergon TL, Lee LFS, Dagit RG, et al. Effects of Urbanization on the Distribution and Abundance of Amphibians and Invasive Species in Southern California Streams. Conservation Biology. 2005 Dec;19(6):1894–907.

63. Riley SPD, Sauvajot RM, Fuller TK, York EC, Kamradt DA, Bromley C, et al. Effects of Urbanization and Habitat Fragmentation on Bobcats and Coyotes in Southern California. Conservation Biology. 2003 Apr;17(2):566–76.

64. Ruas RDB, Costa LMS, Bered F. Urbanization driving changes in plant species and communities – A global view. Global Ecology and Conservation. 2022 Oct;38:e02243.

65. Seto KC, Güneralp B, Hutyra LR. Global forecasts of urban expansion to 2030 and direct impacts on biodiversity and carbon pools. Proceedings of the National Academy of Sciences. 2012 Oct 2;109(40):16083–8.

66. Shirey V, Belitz MW, Barve V, Guralnick R. A complete inventory of North American butterfly occurrence data: narrowing data gaps, but increasing bias. Ecography. 2021 Apr;44(4):537–47.

67. Sodhi NS, Bickford D, Diesmos AC, Lee TM, Koh LP, Brook BW, et al. Measuring the Meltdown: Drivers of Global Amphibian Extinction and Decline. PLoS ONE. 2008 Feb 20;3(2):e1636.

68. Spotswood EN, Beller EE, Grossinger R, Grenier JL, Heller NE, Aronson MFJ. The Biological Deserts Fallacy: Cities in Their Landscapes Contribute More than We Think to Regional Biodiversity. BioScience. 2021 Feb 15;71(2):148–60.

69. Steen VA, Elphick CS, Tingley MW. An evaluation of stringent filtering to improve species distribution models from citizen science data. Diversity and Distributions. 2019 Dec;25(12):1857–69.

70. Steen VA, Tingley MW, Paton PWC, Elphick CS. Spatial thinning and class balancing: Key choices lead to variation in the performance of species distribution models with citizen science data. Methods in Ecology and Evolution. 2021 Feb;12(2):216–26.

71. Szabo JK, Khwaja N, Garnett ST, Butchart SHM. Global Patterns and Drivers of Avian Extinctions at the Species and Subspecies Level. PLoS ONE. 2012 Oct 8;7(10):e47080.

72. Uchida K, Blakey RV, Burger JR, Cooper DS, Niesner CA, Blumstein DT. Urban Biodiversity and the Importance of Scale. Trends in Ecology and Evolution. 2021 Feb;36(2):123–31.

73. Urošević A, Tomović L, Ajtić R, Simović A, Džukić G. Alterations in the reptilian fauna of Serbia: Introduction of exotic and anthropogenic range expansion of native species. Herpetozoa. 2016;28(3/4):115–32.

74. Van Strien AJ, Van Swaay CAM, Termaat T. Opportunistic citizen science data of animal species produce reliable estimates of distribution trends if analysed with occupancy models. Devictor V, editor. Journal of Applied Ecology. 2013 Dec;50(6):1450–8.

75. Veech JA, Small MF, Baccus JT. The effect of habitat on the range expansion of a native and an introduced bird species: Habitat and range expansion. Journal of Biogeography. 2011 Jan;38(1):69–77.

76. Wehtje W. The range expansion of the great-tailed grackle (*Quiscalus mexicanus Gmelin*) in North America since 1880: Range expansion of the great-tailed grackle. Journal of Biogeography. 2003 Oct;30(10):1593–607.

77. West S, Pateman R. Recruiting and Retaining Participants in Citizen Science: What Can Be Learned from the Volunteering Literature? Citizen Science: Theory and Practice. 2016 Dec 31;1(2):15.

78. Wood EM, Esaian S. The importance of street trees to urban avifauna. Ecological Applications. 2020 Oct;30(7):e02149.

79. Wooster EIF, Fleck R, Torpy F, Ramp D, Irga PJ. Urban green roofs promote metropolitan biodiversity: A comparative case study. Building and Environment. 2022 Jan;207:108458.

80. Yang J. Big data and the future of urban ecology: From the concept to results. Science China Earth Sciences. 2020 Oct;63:1443–56.

